# *De novo* mutations mediate phenotypic switching in an opportunistic human lung pathogen

**DOI:** 10.1101/2024.02.06.579193

**Authors:** Alexandra J. Poret, Matthew Schaefers, Christina Merakou, Kathryn E. Mansour, Georgia K. Lagoudas, Ashley R. Cross, Joanna B. Goldberg, Roy Kishony, Ahmet Z. Uluer, Alexander J. McAdam, Paul C. Blainey, Sara O. Vargas, Tami D. Lieberman, Gregory P. Priebe

## Abstract

Bacteria evolving within human hosts encounter selective tradeoffs that render mutations adaptive in one context and deleterious in another. Here, we report that the cystic fibrosis-associated pathogen *Burkholderia dolosa* overcomes in-human selective tradeoffs by acquiring successive point mutations that alternate phenotypes. We sequenced the whole genomes of 931 respiratory isolates from two recently infected patients and an epidemiologically-linked, chronically-infected patient. These isolates are contextualized using 112 historical genomes from the same outbreak strain. Within both newly infected patients, diverse parallel mutations that disrupt O-antigen expression quickly arose, comprising 29% and 63% of their *B. dolosa* communities by 3 years. The selection for loss of O-antigen starkly contrasts with our previous observation of parallel O-antigen-restoring mutations after many years of chronic infection in the historical outbreak. Experimental characterization revealed that O-antigen loss increases uptake in immune cells while decreasing competitiveness in the mouse lung. We propose that the balance of these pressures, and thus whether O-antigen expression is advantageous, depends on tissue localization and infection duration. These results suggest that mutation-driven alternation during infection may be more frequent than appreciated and is underestimated without dense temporal sampling.

## Main

Bacteria colonizing humans are subjected to environmental variations across both space and time. The optimal survival strategy in one condition can be counterproductive in another, creating selective tradeoffs^1–3^. Bacteria employ a range of strategies to navigate tradeoffs, including evolving environment-responsive regulatory elements^4^, acquiring and discarding genetic programs via horizontal gene transfer^5^, and developing sequence motifs prone to high contraction and expansion rates^6^.

The acquisition of *de novo* mutations to toggle a phenotype on and off has traditionally been considered inefficient due to the low likelihood of restoring a pathway’s original function by random mutation^7,8^. However, reversion has occasionally been observed during *in vitro*^9^ and *in vivo* laboratory evolution experiments^10^, consistent with theoretical work suggesting that reversion is likely under conditions of high mutation rate and high population size^11,12^. Yet, concrete in-human evidence of recurrent *de novo* mutation driving phenotypic switching has remained elusive.

Here, we report a bacterial pathogen acquiring successive *de novo* mutations in the same pathway, including reversion of a stop codon, to navigate repeated environmental change within and between hosts. We compare early versus long-term infections with *Burkholderia dolosa*, which causes chronic lung infections in people with cystic fibrosis (CF)^13^, using 931 whole genomes isolated from newly acquired and long-term chronic infections. Our phylogenetic analysis reveals that *B. dolosa* switches O-antigen expression on and off via *de novo* mutation, that O-antigen expression is more advantageous in later infection stages, and that O-antigen loss is adaptive at early infection stages. *In vivo* experiments in mice demonstrate that while O-antigen expression is beneficial for *B. dolosa* survival in the lung, it is detrimental in the spleen. Together, our results show how *B. dolosa* repeatedly mutates the same pathway to navigate a selective landscape that changes over the course of chronic infection.

### Selection for O-antigen expression during long-term infection

In the 1990s, an outbreak of *B. dolosa* spread among CF patients in the Boston area^14^. We previously studied the genomic evolution of *B. dolosa* during this outbreak using a longitudinal collection of 112 isolates^15^ and culture-based metagenomics^16^ from sputum samples collected after 7-9 years of chronic infection. These studies revealed genes under parallel evolution and pointed to key survival strategies of bacteria in the CF lung^15,16^.

One of the most notable signatures for adaptation during this outbreak was seen in the pathway encoding for *B. dolosa*’s O-antigen, the sugar-chain decorating the lipopolysaccharide (LPS)^15^. Surprisingly, phylogenetic analyses indicated that this outbreak was initiated by a strain carrying a stop codon in *wbaD* (AK34_RS24375, previously annotated as BDAG_02317), a glycosyltransferase gene essential for the expression of O-antigen. This stop codon underwent at least 10 independent reversions, reinstating O-antigen expression. As these chronic infections persisted, the proportion of isolates expressing O-antigen increased steadily. Variation in O-antigen expression and length has also been widely documented in other *Burkholderia* species that chronically infect the CF lung^17–20^.

The ability of a single mutation to restore function suggests this gene was pseudogenized only in the recent past. Moreover, phylogenetic data suggest that initial infections during this outbreak were established with O-antigen-absent strains^15^. These observations raise the intriguing possibility that O-antigen presence is disadvantageous during transmission or early infection, despite its advantage during later stages of disease.

### New infection cluster was established by *B. dolosa* genotypes expressing O-antigen

With the guiding hypothesis that O-antigen expression may be a liability during transmission or early stages of infection, our lens focused on a new *B. dolosa* transmission cluster in the Boston vicinity after 9 years of no new cases. This cluster comprised 3 adults: Subject J (the suspected index patient), Subject Q (co-worker of Patient J), and Subject R (sibling of Patient Q) (**Fig. 1**). Patient J was known to be infected with *B. dolosa* as part of the historical outbreak. Siblings Q and R, the new patients, lived together and were not followed at the hospital of the historical outbreak.

**Figure 1:**
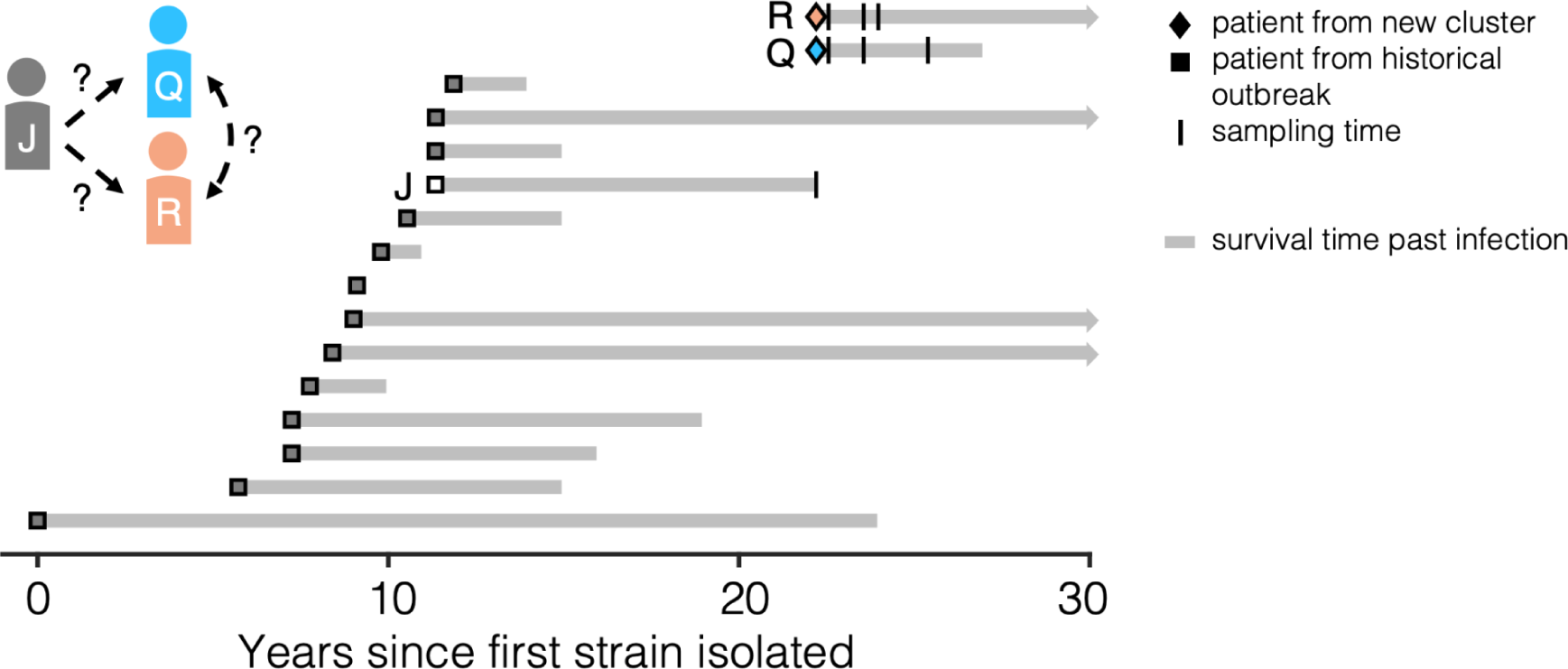
A new suspected *Burkholderia dolosa* transmission cluster among 3 adults with cystic fibrosis with connection to a historical outbreak. Two siblings with cystic fibrosis (Patients Q and R) were recently infected with *B. dolosa* and suspected to be part of a three-person transmission cluster (with Patient J). Patient J was previously infected as part of a historical outbreak that infected 39 people, 14 of whom were profiled in a previous study^15^ and whose isolates are analyzed here. About 10 years elapsed between the last known transmission in that outbreak and the latest cluster. Here, we sequenced a total of 122 isolates cultured from the sputum of the new Patients Q and R over multiple timepoints (vertical bars). We also acquired 650 isolates from the lungs, lymph nodes, and spleen during autopsy of Patient J and 159 from blood obtained a week prior. Gray bars indicate survival time past infection; arrowheads on the far right indicate that the patient was alive as of January 2024.

To test the hypothesis that O-antigen expression is disadvantageous during transmission or early infection, we characterized the *B. dolosa* populations in this new Boston-area transmission cluster. We collected three sputum samples from Patients Q and R between 4 and 38 months after first diagnosis and sequenced the genomes of 5-24 single-colony *B. dolosa* isolates from each sample (122 total; **Fig. S1**). The death of Patient J due to influenza and cepacia syndrome in the same year enabled us to study Patient J’s *B. dolosa* population in great detail: we sequenced 650 isolates collected during autopsy and 159 isolates from multiple blood cultures drawn in the week prior to death (**Table S1**). We also reanalyzed 10 genomes from Patient J from a prior study, reconstructing 16 years of within-person evolution^15^.

Evolutionary reconstruction indicated that Patient J was the infection source for Patients Q and R. Across a genome-wide phylogeny encompassing 927 newly sequenced and 112 previously-published isolates taken a decade prior from 14 patients, Patient Q’s & R’s isolates are nested within a clade of Patient J’s isolates. This topology strongly supports the epidemiological association between Patient J and the new patients (**Fig. 2a, Fig. S2, Fig. S3**). Yet, the phylogenetic structure does not resolve transmission between the siblings. Neither siblings’ isolates are monophyletically nested within another’s isolates, leaving room for multiple hypotheses: back-and-forth transmission between siblings, transmission of the same multiple genotypes from J to both siblings, or parallel evolution creating the same exact nucleotide mutation in both siblings. Patients Q and R may have been more likely to experience complex transmission because of their cohabitation. Schematics of hypothetical transmission scenarios are in **Fig. 2b**; each of these requires the founding of at least one patient’s population by two or more independent genotypes (arrows in **Fig. 2b)**, suggesting that *B. dolosa* infections in CF don’t always hinge on a single cell transmission.

**Figure 2:**
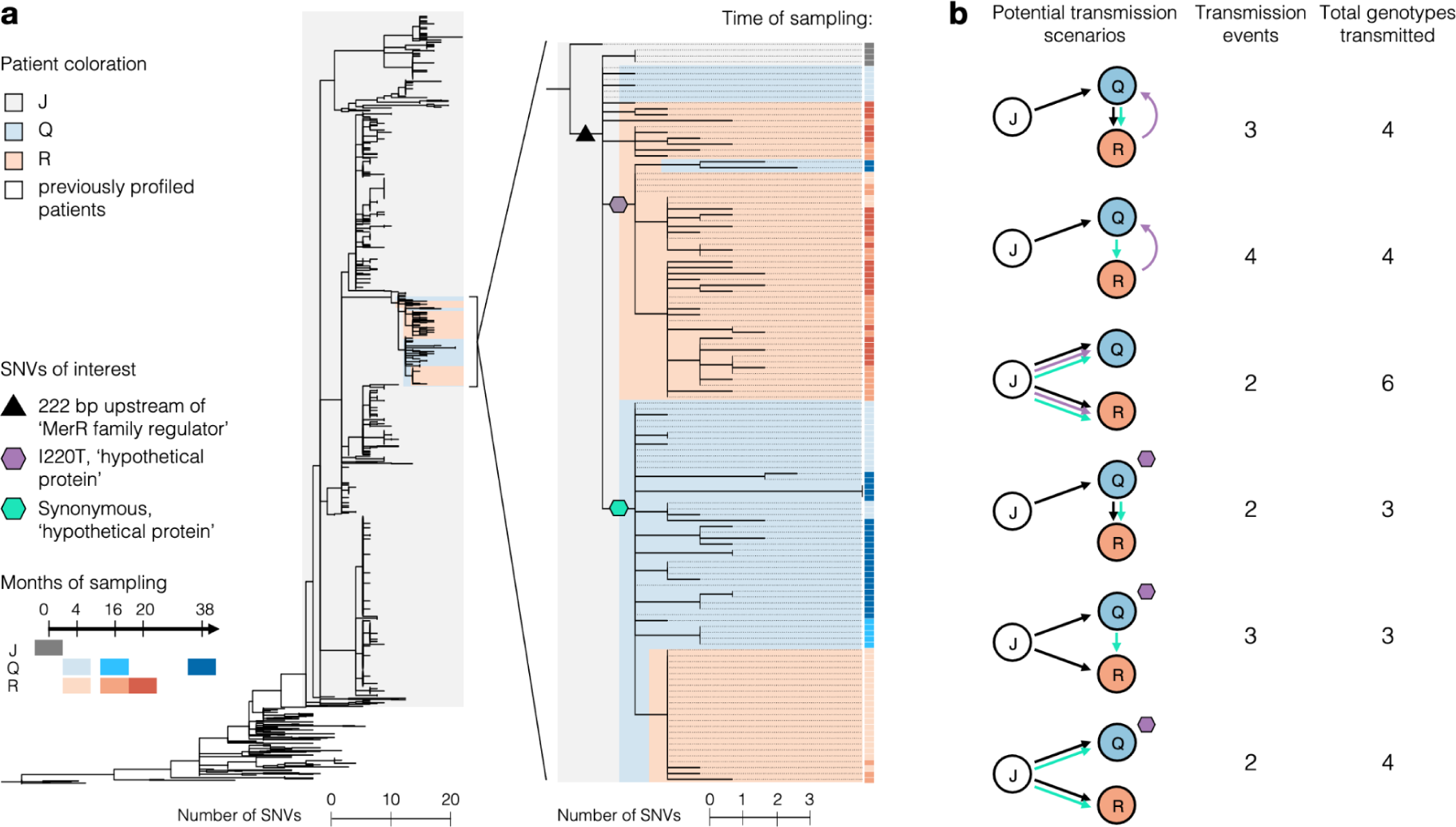
In early infection establishment, multiple *B. dolosa* clones are transmitted between patients. (a) A maximum parsimony SNV phylogeny was built from whole-genome sequencing of 805 *B. dolosa* isolates from autopsy samples of the suspected index case (Patient J), 122 from serial sputum samples of Patients Q and R, and 112 previously published sequences of isolates from Patient J and 13 other patients a decade prior. All isolates from Patient Q and R are descended from a subset of Patient J’s diversity; a high resolution phylogeny of this clade (Methods) is shown to the right. The time of sampling of each isolate is indicated by color in the rightmost vertical bar, clades within the tree are shaded by patient, and SNVs of interest for transmission inference are indicated by colored shapes. (b) Potential transmission scenarios between Patients J, Q, and R are diagrammed, showing the need for either more than two transmission events or parallel nucleotide evolution to explain observed diversity. Arrows point in the direction of *B. dolosa* transfer and are colored by the SNV in panel (a) that precedes a transmission.

Despite this complexity, we can discern the genotype of the most recent common ancestor (MRCA) of these closely related strains -- and, by proxy, the O-antigen phenotype. Our prior observations^15^ led us to expect an O-antigen-deficient MRCA, containing the truncated *wbaD*. This variant was present in just 1.4% of isolates in Patient J at the time of death (**Fig. S4**), making this scenario possible but unlikely. Contrary to our selection-based prediction, Patient Q’s and R’s MRCAs encode for the full length *wbaD*, and the earliest isolates from these subjects expressed O-antigen. These predicted phenotypes were confirmed experimentally (**Fig. S5**). This observation demonstrates that O-antigen presence does not prevent transmission and thus demands another explanation for the absence of O-antigen in the early stages of the larger historical outbreak.

### Selection for O-antigen expression during short-term infection

Soon after establishment of this O-antigen-expressing-strain in Patients Q and R, diverse *de novo* mutations quickly arose, converging on the O-antigen pathway. Across both patients, a total of 18 independent, parallel mutations were observed in this pathway (**Fig. 3a**). This concentration of mutations is statistically significant when compared to a random genomic distribution (**Fig. 3b**; P < .01, binomial test) and when compared to mutations accumulated in Patient J over a decade of infection and detected at the time of autopsy (**Fig. 3b**; P < .01, binomial proportion test). These results support a model in which disruptions in O-antigen expression are strongly selected for during early *B. dolosa* infection.

**Figure 3:**
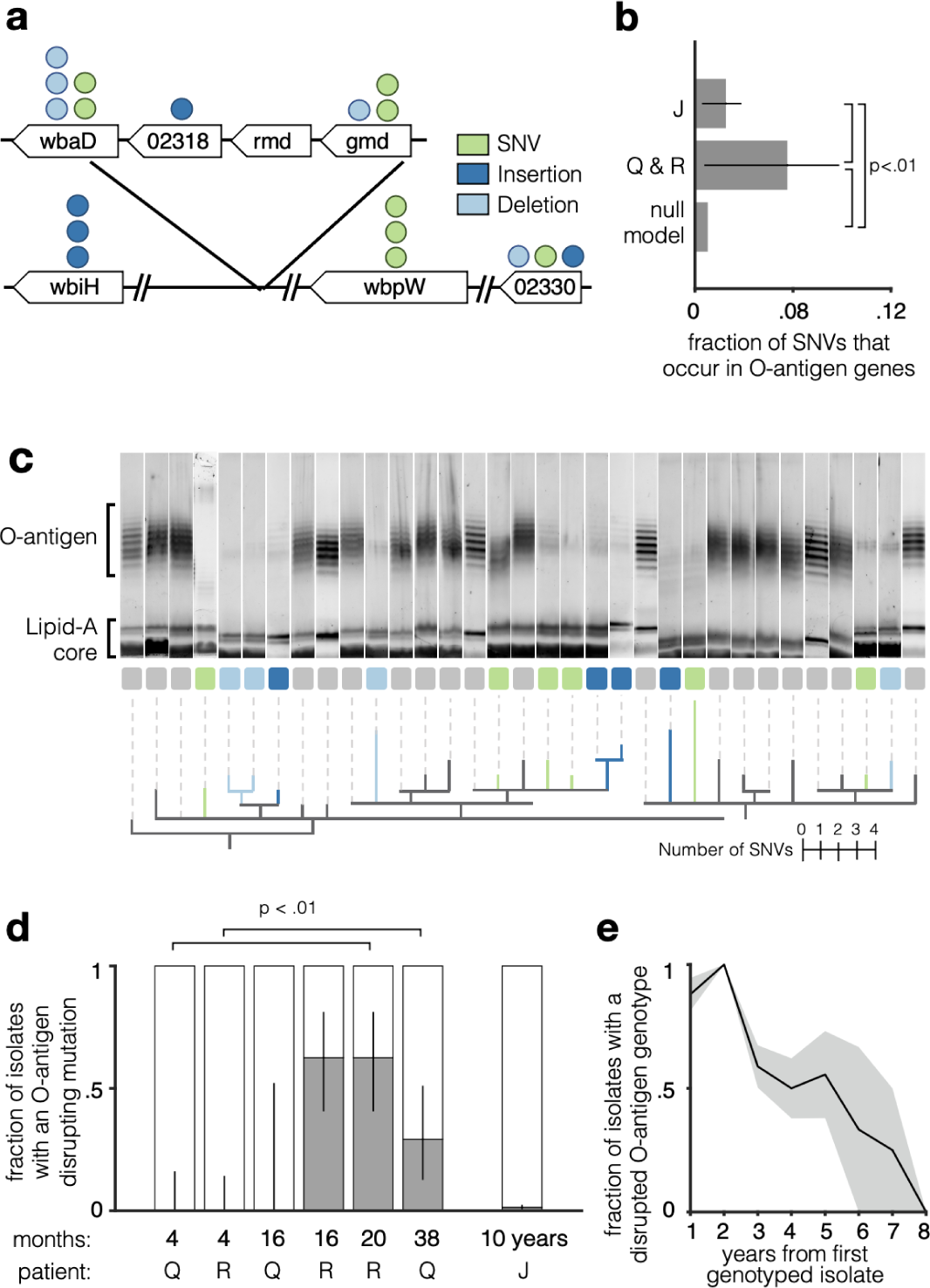
*B. dolosa* lipopolysaccharide (LPS) O-antigen expression is reduced in early CF lung infection by *de novo* mutation, contrasting with trends in longer-term colonization. (a) Parallel mutations in genes suspected to be involved in O-antigen synthesis are represented by circles. Each isolate contained at most one such mutation. Mutations are colored by the type of mutation and gene icon length is proportional to gene size. (b) The proportion of observed SNVs that fall in the O-antigen synthesis is significantly greater for Patients Q and R (recently infected) than for Patient J (infected for nearly a decade; P < .01 binomial proportion test). Both sets of mutations occur at a greater frequency than expected by random mutation across the O-antigen pathway (P < .01, binomial test, see Methods). Error bars indicate 95% confidence intervals. (c) Selected isolates from Patient Q and R, as well as near-isogenic samples from Patient J, were profiled for O-antigen banding pattern. Isolates are ordered by a phylogeny of SNVs and O-antigen-affecting indels (see Methods) and branches are colored by the occurrence of an O-antigen-affecting SNV, insertion, or deletion. Of 14 isolates with an O-antigen-affecting mutation, 13 show a reduction of O-antigen expression. (d) The proportion of O-antigen-disrupted isolates is significantly increased from the first to last sampled time point for both Patients Q and R (P < .01, binomial proportion test), but negligible in patient J after 10 years. Lines represented with 95% confidence intervals. (e) In a reanalysis of 112 isolate genomes from the original *B. dolosa*^15^ outbreak collected over a greater duration of infection, we identify a trend during extended chronic infection that contrasts with early infection. See Fig. S6 for more details. The shaded region indicates standard error of the mean.

To probe the impact of these mutations on O-antigen expression, we stained for the O-antigen in 14 isolates with distinct mutations in this pathway, alongside 19 control isolates within the same clade without any mutations predicted to affect O-antigen synthesis. While all control isolates from Patients Q & R displayed the O-antigen, 13 out of the 14 isolates with a mutation in the O-antigen pathway (93%) showed reduced O-antigen expression -- indicating that most of the identified mutations impacted O-antigen synthesis (**Fig. 3c**). These results provide confidence in our genotype-to-phenotype predictions.

Combining this phenotype map with the entirety of the genotypic data, we observed a strong temporal change in the prevalence of O-antigen presentation. At four months post-infection, all *B. dolosa* isolates from Patients Q & R bore O-antigen-expressing genotypes. Yet, by 20 and 38 months, 29% of Q’s and 63% of R’s isolates had acquired a mutation (either SNV or indel) that disrupted O-antigen expression (**Fig. 3d**; P < .01, binomial proportion test). This signal for early O-antigen loss contrasts with the prior observation of selection for O-antigen late during chronic infection (**Fig. 3e**) and indicates an advantage for O-antigen disruption specifically during the first years of chronic infection.

### Diverse O-antigen chain lengths coexist over a decade of chronic infection

To better understand O-antigen evolution in late infection, we examined the emergence of O-antigen-affecting mutations across Patient J’s 809 isolates and phenotyped representative clones. This revealed four distinct O-antigen phenotype classes, each paired with a distinct genotypic signature: (1) absent, without any expression of O-antigen due to the stop codon in *wbaD*, (2) long-chain, found in isolates with the restored *wbaD* and no other mutations impacting O-antigen expression; (3) medium-chain, found in isolates with a restored *wbaD* and a 3 base pair insertion in BDAG_02328 (AK34_RS24445), encoding a different glycosyltransferase; and (4) short-chain, found in isolates with the previous two mutations plus a mutation in a third glycosyltransferase BDAG_02321 (AK34_RS24395) (**Fig. S4, S5**). Genotypes corresponding to all 4 of these phenotypes were also found among isolates from Patient J collected 5-11 years earlier **(Table S2).** To assess the generality of long-term coexistence, we considered isolates from other patients in the historical *B. dolosa* outbreak^15^. Indeed, we find multiple patient timeseries for which O-antigen-absent strains were recovered years after an O-antigen-intact strain was observed (in 4 of 13 cases). In addition, mutations that shorten the O-antigen chain were also observed in 3 of 13 patients (**Fig. S6, S7**). Together, these findings indicate that diverse O-antigen phenotypes can coexist within an individual for years.

Within Patient J’s lungs, the O-antigen absent phenotype was not enriched at any lung lobe, though we noted a modest difference in other O-antigen phenotypes across lobes (**Fig. S8, S9, S10**). Analysis of *B. dolosa* spreading within the lung using mutational distances^21,22^ showed only modest signal for spatial stratification. This lack of a strong spatial signal might be attributed to a breakdown of lung integrity following severe influenza and mechanical ventilation. Alternatively, given the relatively large postmortem lung tissue sizes (∼1cm^2^), each sample might encompass multiple microenvironments (e.g. intracellular vs. extracellular, epithelial tissue vs. immune cells), each potentially favoring alternative O-antigen phenotypes.

### Tradeoffs in murine spleen and lung colonization

What, mechanistically, can explain the time-based inversion of selection on O-antigen presentation? Expression of O-antigen is known to provide resistance to serum complement^23,24^, antibiotics^17^, and antimicrobial peptides^25^, and is therefore expected to outcompete O-antigen-disrupted strains in most conditions. However, given its recurrent loss during early infection, O-antigen expression must be selected against in some environmental conditions.

To discern a potential advantage for *B. dolosa* O-antigen disruption across body compartments, we utilized a murine pneumonia model (Methods). We selected 3 independent pairs of strains that differ in O-antigen presentation, each containing an O-antigen-disrupted strain and near-isogenic strain with an intact LPS (**Fig. S11,** Methods). For each pair, we performed *in vivo* competitions by labeling the O-antigen-intact strain with a transposon that constitutively expresses LacZ. This labeled strain was mixed with its un-labeled O-antigen-disrupted partner and used to intranasally infect mice (**Fig. 4a**). After 7 days, the ratio of O-disrupted to O-antigen-intact strains was significantly greater in the spleen than in the lungs, for all pairs (**Fig. 4b**; 1.6 to 2.4-fold, paired T-test, P < .002). We also performed label-swap experiments for each pair, where the O-antigen-disrupted strain was marked with LacZ, and saw similar findings (**Fig. S12**). Slight variations were seen among pairs of isolates, potentially due to differences in the O-antigen-disrupting mutation or other genomic variations. Nevertheless, the organ-based tradeoff was consistent across isolate pairs and label-swaps.

**Figure 4:**
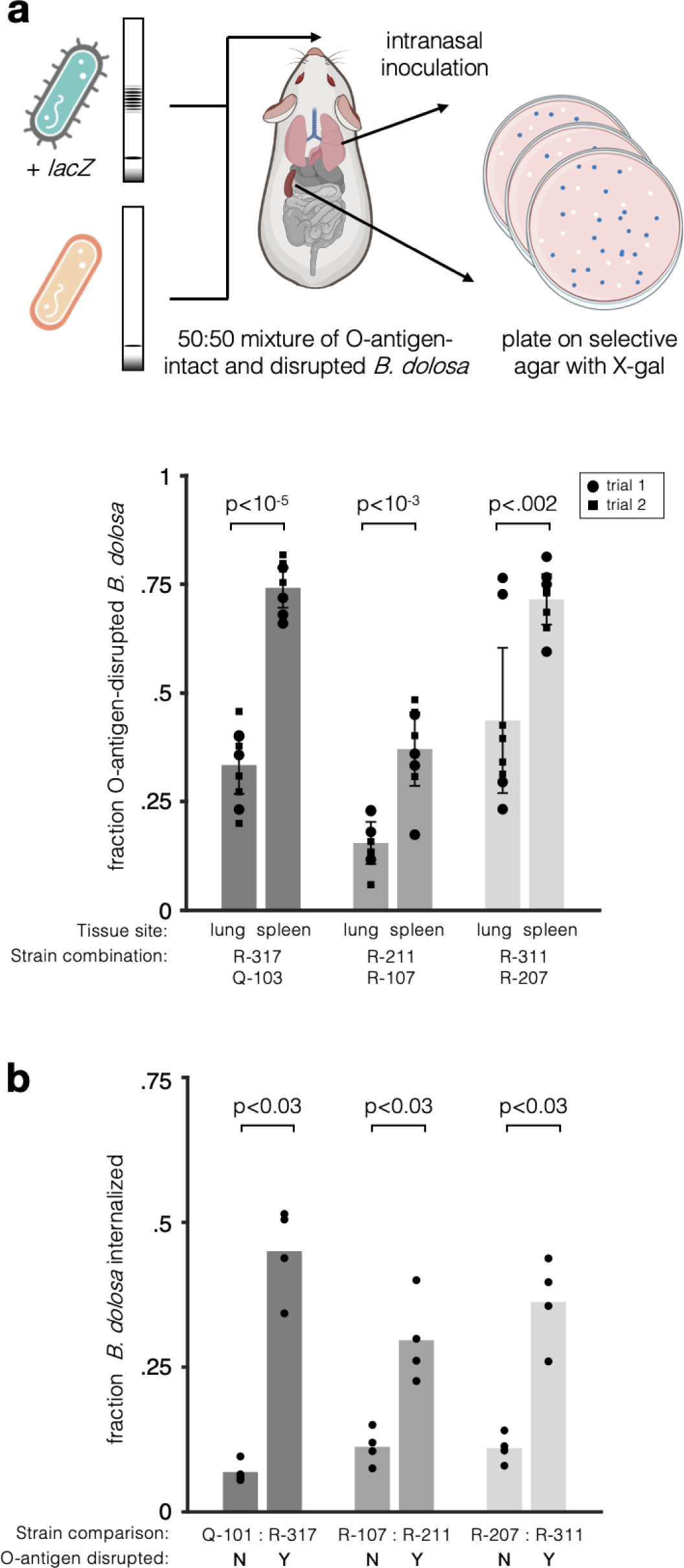
Disruption of *B. dolosa* LPS O-antigen expression leads to tradeoffs in lung versus spleen infection *in vivo*. (a) Mice were infected with a 50:50 mixture of O-antigen-intact and disrupted phenotypes, with one *B. dolosa* strain marked with a *lacZ* cassette conferring blue colony color on X-gal-containing *Bcc*-selective agar. Lungs and spleens were excised to determine the bacterial load of O-antigen-intact and O-antigen-disrupted strains. Two trials of this experiment were performed for each of three near-isogenic pairs. The fraction of O-antigen-disrupted bacteria in mouse spleens and lungs is shown with error bars representing 95% confidence intervals. The O-antigen-disrupted to O-antigen-intact ratio is higher in spleens relative to lungs for all pairs (P < .002, paired T-test). (b) Macrophages were infected for 2 hours with near-isogenic *B. dolosa* strains with and without an O-antigen-disrupting mutation and then incubated for 2 hours with kanamycin, which kills extracellular bacteria. The number of intracellular bacteria is compared to the total bacteria obtained from an identical culture without kanamycin treatment, revealing an advantage for O-antigen-disrupted bacteria within macrophages (P < .03, Wilcoxon rank sum).

Given the prominence of O-antigen-disrupted genotypes in the spleen, we hypothesized that O-antigen might affect *B. dolosa*’s ability to invade and persist within phagocytic cells^26^. The intracellular niche is known to be critical to establishment of infection in the broader *Burkholderia cepacia* complex (Bcc)^27,28^, and O-antigen negativity has previously been shown to increase internalization of other Bcc species^29^. We therefore infected macrophages with O-antigen-intact and disrupted strains. In line with our expectation, the number of intracellular bacteria was 1.7 - 8.2 fold higher for O-antigen-disrupted strains after two hours of macrophage infection (**Fig. 4b, Fig. S13**, P < .03 for 8/9 comparisons, Wilcoxen rank sum). Combined with the mouse data, this result confirms the superiority of the O-antigen-disrupted strains within immune cells.

Together, these results suggest that O-antigen disruption is selected for during the first years of infection in the CF lung because of its advantage in growth or infiltration of phagocytic cells, and that this is superseded by changing environmental pressures during the course of longer chronic infection, perhaps related to changing oxygen availability with increasing lung damage^30^, increased antibiotic pressures, or to adaptive immune responses^31^ (**Fig. 5**). However, we cannot rule out the possibility that other mechanistic forces, such as the higher propensity of O-antigen-disrupted strains to adhere to host cells^29^, could also be influencing the selective landscape.

**Figure 5:**
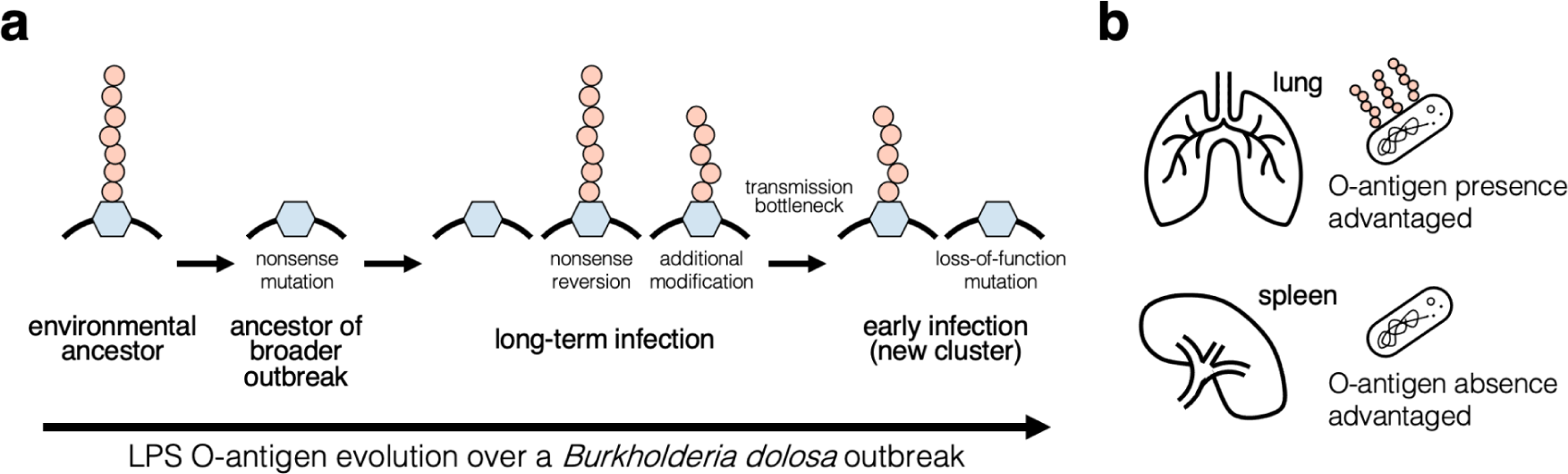
Navigation of selective tradeoffs on O-antigen presentation by *de novo* mutation. (a) Inferred natural history of *B. dolosa* LPS O-antigen presentation during an outbreak in people with CF. The initial *B. dolosa* outbreak was initiated by a strain lacking O-antigen due to a premature stop codon in a glycosyltransferase gene. This premature stop mutation likely occurred shortly before or during outbreak initiation, as experimental reversion successfully restores O-antigen presentation^15^. As the outbreak spread and persisted in chronic infections, independent reversions restored *B. dolosa*’s O-antigen in multiple patients. Within each patient, strains with variable O-antigen presentation were recoverable and coexisted for years. Two patients recently infected by a survivor of this outbreak were initially colonized by strains expressing O-antigen. Remarkably, O-antigen-disrupted phenotypes re-emerged independently within each of these patients via a variety of *de novo* mutations in the O-antigen pathway. (b) Our *in vivo* experimental results suggest that O-antigen presence is advantageous in the lung while O-antigen absence is favored in the spleen. Combined with the historical observations, these results suggest that survival in the spleen or immune cells may be important during the early years of chronic *B. dolosa* lung infection.

## Discussion

In this study, we show that *B. dolosa* adapts and diversifies in response to shifting environmental pressures on LPS O-antigen presentation within individual people. O-antigen expression is selected for within immune cells and in early infection, while its absence is selected for within the lung and during late infection. In response to these tradeoffs, *B. dolosa* cells switch O-antigen expression on and off through a seemingly permanent solution -- mutation -- only to revert to the previous phenotype through additional mutation.

Our results illuminate previous observations of phenotypic variation in O-antigen presentation among members of the *Burkholderia cepacia* complex (Bcc). Conflicting observations of loss^19,20,32^ and acquisition of O-antigen in *Burkholderia cepacia complex* species^15,19^ can be rationalized when viewed through the lens of context-dependent mutations. This clarity was enabled by a combination of deep and longitudinal population sequencing, phenotypic profiling, and *in vitro* and *in vivo* models of infection.

To our knowledge, this is the most direct observation of repeated phenotypic switching via conventional mutation outside of a laboratory setting. Even though prior *in vitro* studies^33^ and controlled mouse experiments^10^ observed mutation-mediated switching, doubt has been cast on the relevance of these phenomena in natural settings^7,8^. Instead, a change in selective pressures after the acquisition of a pleiotropic point mutation is considered more likely to select for one or more of the following scenarios: the emergence of a compensatory mutation^12^; the migration of cells with ancestral phenotypes^34^; or the resurgence of a persistent, low-frequency ancestral genotype^35,36^. Mutation-driven switching is commonly relegated only to genomic elements with elevated mutation rates (e.g. simple sequence repeats^37–39)^. Our work challenges this expectation by presenting an *in situ* case of bacteria undergoing mutation-mediated phenotypic switching and calls for theoretical models that consider mutation-mediated reversion.

Mutation-mediated switching and reversion may well be common among bacteria with within-host population sizes exceeding the inverse of the per-nucleotide mutation rate (∼10^-9^ mutations/generation for most species^40^), including other CF lung pathogens and commensal members of the gut microbiome^2,11,41^. Mutation-driven adaptations could be considered less likely for commensal and other organisms with prolonged tenure in mammalian environments which have had ample time to evolve more complex regulatory mechanisms^42,43^. However, recent observations suggest that mutation-mediated phenotypic switching might be common even for species with ample time to evolve gene regulation: *E. coli* undergoes mutation-mediated switching in laboratory mice^10^ and *B. fragilis’* in-human adaptive mutations target conserved genes^44^, despite these same genes possessing complex regulatory mechanisms^38^.

Altogether, our work suggests that the speed, dynamism, and consequences of bacterial adaptations to the ever-fluctuating human environment^33,45–47^ may be underappreciated. It has been speculated that within-host adaptations to these changing fitness landscapes represent ‘dead-ends’ that do not transmit to others^9,34^. However, the observation of nucleotide-level and phenotype-level reversions seen here suggest that additional studies with deep intrapopulation sampling along a transmission chain are needed to fully assess the scope of mutation-mediated phenotypic variability. If mutation-mediated phenotypic switching is indeed widespread, the extent to which within-human bacterial evolution drives phenotypic change may be vastly underestimated.

## Methods

### Study cohort, isolate sampling, and genome sequencing

Starting in 1998, 39 cystic fibrosis patients were infected with *Burkholderia dolosa* during an outbreak in Boston Children’s Hospital, Boston, MA. No new *B. dolosa* infections were reported in the Boston area between 2005 and 2013, and the outbreak was presumed to have ended. In 2014, a pair of siblings (Patient Q and Patient R) were confirmed to have newly acquired *B. dolosa* infections, and a likely index patient was identified. The suspected index patient of this new cluster was infected during the known previous outbreak, and this patient’s *B. dolosa* population was described in two previous studies: Patient J in Lieberman and Michel et al 2011^15^ and Patient 2 in Lieberman et al 2014^16^; referred to as Patient J in this study).

Sputum samples were obtained during normal care under IRB approved protocol 05-02-014R (with written informed consent) at Boston Children’s Hospital (BCH) from Patients Q and R at 3 timepoints following initial diagnosis: 4 months, 16 months, and either 20 (Patient R) or 38 months (Patient Q) (**Fig. 1; Table S1**). Sputum samples were diluted in 1% proteose peptone and then plated on oxidation/fermentation-polymyxin-bacitracin-lactose (OFPBL) media (BD Biosciences) to select single *B. dolosa* colonies. From every sample, up to 24 colonies were randomly picked and frozen in 15% glycerol; in the event of low CFU density, we sampled all present colonies. This resulted in 123 total isolates, with 5-24 collected per patient timepoint.

Patient J died from cepacia syndrome developed during hospitalization for influenza shortly after identification of the new *B. dolosa* cases. Blood cultures grew *B. dolosa* over the 7 days prior to death. Blood culture bottles found to be growing *B. dolosa* were obtained under the same IRB protocol as above. Contents of blood culture bottles were diluted in 1% proteose peptone and then plated on OFPBL plates for colony picking. We collected 170 total blood colonies, ranging from 16-29 isolates per time point. During Patient J’s autopsy, we obtained lung (599), spleen (47), and lymph node (48) isolates from 38 different sites (**Fig. S8**) under the same IRB protocol as above. Tissue samples were taken using sterile technique, homogenized, and frozen in glycerol as described in Chung et al^21^. Serial dilutions of lung, spleen, and lymph node tissue were plated on OFPBL media. Up to 24 *B. dolosa* colonies were grown from each sample for 24h in Luria broth (LB) and then frozen in glycerol.

Bacteria were cultured to mid-log in LB for DNA extraction, library preparation, and DNA sequencing. The standard Illumina Nextera protocol was used in a modified method for library construction using a microfluidic device that enabled parallel and resource-efficient DNA library preparation. Microfluidic methods were based on Kim et al.^48^ and further updated, as described in a forthcoming publication. The resultant libraries were sequenced to a median depth of 87.7 on an Illumina HiSeq X Ten with 2 x 150 bp dual indexed reads.

### Mutation detection and phylogenetic inference

Adaptors were trimmed by cutadapt v1.18^49^ and then filtered via sickle v1.33^50^ (filters: “-q 20 -l 50 -x -n”). Filtered reads were aligned by bowtie2^46^ (v2.2.6; bowtie2 -X 2000 --no-mixed --dovetail --very-sensitive --n-ceil 0,0.01) to the reference genome AU0158, a *Burkholderia dolosa* sample isolated from the index patient of the original Boston outbreak five years into infection (Genbank: GCA_000959505.1). SAMtools v1.5^51^, mpileup (-q30 -t SP -d3000), bcftools call (-c), and bcftools view (-v snps -q.75) was used to create consensus sequences and call candidate single nucleotide variants (SNVs) across all isolates. All code used to call these commands was aggregated in a MATLAB v2015b and run on the high performance computing cluster, “Commonwealth Computational Cloud for Data Driven Biology” (C3DDB).

Candidate SNVs and samples were filtered based on coverage, nucleotide identity, and SAMtools produced FQ score in MATLAB v2021a according to the below filters. Samples with a mean candidate SNV coverage of <12x across candidate positions were discarded (17/987 samples). SNV positions were removed if they had a median depth of coverage across samples of < 40x or if more than 20% of sequence calls across samples were marked as ambiguous, where ambiguity is defined as having a major allele frequency of less than .9, a per-strand coverage of less than 12x, or a FQ score below 0 in a given sample.

Once a list of 541 trusted loci were obtained, each sample’s nucleotide call at a given position was defined as the major allele across reads, or labeled as N if the major allele frequency was <.8 or had no reads (**Table S3, Table S4, Table S5**). Any sample with more than 10 trusted positions called as N was removed from further analysis (39/987 samples). In total, 931 samples passed this filtering protocol. The resulting allele calls were used to construct a maximum-parsimony phylogenetic tree of Patients J, Q, and R using DNAPars (Phylip v3.69)^52^. The tree was rooted with the reference genome AU0158 and can be seen in **Fig. S2**. We also identified short insertions and deletions (indels) in and nearby genes affecting the O-antigen presentation (**Table S6**). Indels are difficult to call with high fidelity, and both false-positive and false-negative mutation calls can impact phylogenetic inference. We used Breseq v0.30.0^53^ and its associated program gdtools to identify indels relative to the reference genome AU0158. Variants around homopolymers and variants for which >.05 of samples were called as unknown or deleted by Breseq were discarded. All remaining indels within 100,000 base pairs of an O-antigen-affecting gene as defined by KEGG’s gene ontology^54^ were queried via PaperBlast^55^ to deduce whether a mutation in that gene could impact B. dolosa’s O-antigen (KEGG: 00541, **Table S7, Table S8, Table S9**). After curating all indels that could plausibly impact O-antigen presentation, we created a second maximum-parsimony phylogenetic tree of the new infection cluster, including one additional SNV for each indel in the input file to DNApars. The inclusion of these indels proved useful in understanding the spread of O-antigen-affecting mutations across the phylogeny. An indel-inclusive tree is shown in **Fig. S4**; a subset of this tree is drawn in **Fig. 3d**.

The list of genes used for O-antigen pathway enrichment analyses was assembled by combining all genes listed in KEGG’s “O-antigen nucleotide sugar biosynthesis” pathway (00541) with all genes between BDAG_02299 (AK34_RS24280) and BDAG_02331 (AK34_RS24460), a mutational hotspot seen in this dataset which contains nine glycosyltransferase genes and other enzymes with putative roles in sugar-modification. The upstream cutoff for O-antigen-affecting genes was conservatively defined as the start of a urease gene cassette (BDAG_02298/AK34_RS24275) and the downstream cutoff was placed before gene BDAG_02332/AK34_RS24465, which precedes a series of genes associated in KEGG with “purine metabolism” (pathway 00230) and containing homology to pyr-(“pyrimidine biosynthetic”)-family proteins.

### Reanalysis of 112 *B. dolosa* isolates from a past outbreak

We created a phylogeny that combines the 112 previously-sequenced isolates from Lieberman and Michel et al.^15^ with the 987 new isolates from Patients J, Q and R. All of the 112 historical isolates were processed using the same procedure described above, with the following exceptions: (1) we used looser sickle v1.33 filters to account for differences in sequencing quality (“-q 0 -l 0 -x -n”) (2) reads were aligned to reference genome AU0158 using bowtie2 using settings modified for single reads ( --phred64 --sensitive --n-ceil 0,0.01). Using the aforementioned SAMtools v1.5, bcftools call, and bcftools view settings, we created consensus sequences and called candidate SNVs across all 1099 total isolates.

To identify loci of interest, we analyzed each set of samples separately; this minimizes the impact of coverage and error bias between sequencing types. We use the same procedure described above to reanalyze and filter Patient J, Q, and R’s samples. We similarly identify SNVs among Lieberman and Michel et al.’s historical samples, with the following modifications: (1) Each allele call at a given position was defined as ambiguous if it had a major allele frequency of less than .9 or coverage of less than 20 reads; (2) No samples were excluded based on coverage; (3) Positions with a median coverage across samples below 20x were removed.

We combine these two candidate loci sets into one trusted list, and define each sample’s nucleotide call at a given position as above. New isolates from Patients J, Q, and R with greater than 10 trusted positions called as N were removed from further analysis. These quality control steps resulted in 927 filter-passing isolates; no such cutoff was implemented for historical samples (which overall had lower sequencing depth and quality). A phylogenetic tree was constructed as above to better understand transmission within the scope of the broader *B. dolosa* epidemic (**Fig. 2A**).

Additionally, we ran Breseq (--predict-polymorphisms) on these 112 historical samples and searched for O-antigen-affecting mutations using the process defined above (**Table S2, Table S10**). Since these isolates are sequenced to a lower depth than those from patients Q, R, and J, we utilize the --predict-polymorphisms setting to increase sensitivity. As such we do not construct a new, indel-aware phylogeny that combines historical samples with our new, differently-processed sequenced data.

### O-antigen phenotyping and quantification

Frozen stocks of *B. dolosa* were used to inoculate 1-5 ml of overnight cultures. These were normalized to an OD_600_ of .25, pelletized using a microcentrifuge at 10,600x *g* for 10 minutes, and stored at −20°C. The next day pellets were thawed and LPS was extracted according to Davis and Goldberg^56^, stopping after the addition of proteinase K (step six) to store overnight at −20°C. Subsequently, 15 ul of each sample were loaded into each well of a 12% Mini-PROTEAN TGX gel (Bio-Rad) along with a CandyCane glycoprotein ladder (Thermofisher). We additionally loaded the same two *B. dolosa* samples with different LPS phenotypes (Q-103; medium and R-221: disrupted) into each gel to assess phenotype repeatability. We run SDS-PAGE on each gel and stain the separated LPS bands using Pro-Q Emerald 300 Lipopolysaccharide Gel Stain Kit (Thermofisher) according to the manufacturer’s instructions, modifying the initial fixation step to be repeated twice and each washing step three times. Images were taken using 300 nm light in a Syngene G:Box Mini-9 imager. The resulting images were processed to quantify the expression and size of the O-antigen.

### Kanamycin exclusion assays to measure *B. dolosa* phagocytosis and survival in macrophages

To understand how different O-antigen phenotypes survived within macrophages, we used a kanamycin exclusion assay from Schaefers et al^57^. Human THP-1 monocytes obtained from ATCC were grown in RPMI 1640 medium (Gibco) containing 2 mM L-glutamine, 10 mM HEPES, 1 mM sodium pyruvate, 4,500 mg/liter glucose, 1,500 mg/liter sodium bicarbonate supplemented with 10% heat-inactivated fetal bovine serum (Gibco), and 0.05 mM 2-mercaptoethanol at 37°C with 5% CO_2_. For differentiation into macrophages, THP-1 cells were diluted to 7 × 10^5^ cells/mL in fresh media and treated with 200 nM phorbol 12-myristate 13-acetate (PMA); 1 mL/well of cells were plated into 24-well plates. Cells were incubated for 3 days as described above. Cells were washed 2x with dPBS (with calcium and magnesium), and RPMI 1640 medium, lacking PMA, was replaced before being incubated in similar conditions for an additional 24 hours.

Cultures of each *B. dolosa* strain used in this analysis, R-317, Q-103, R-211, R-107, R-311, R-207, as well as Q-101, R-318, and R-221, were concurrently streaked out onto OFPBL petri dishes and then incubated overnight in LB while shaking at 200 rpm at 37°C. After subculturing and dilution, log-phase cultures of each bacteria were washed in RPMI three times and ∼2 × 10^6^ CFU were added to each well containing THP-1 macrophages (multiplicity of infection [MOI] of ∼10:1). To synchronize infection, plates were spun at 500 × g for 5 minutes and then incubated for 2 hours at 37°C with 5% CO_2_. After incubation, the total number of bacteria in the wells were determined by adding 100 μL of 10% Triton X-100 lysis buffer to each well with a final concentration of 1%. Having burst the macrophages with Triton, we serially diluted and plated the resultant mixtures on tryptic soy agar (TSA) plates to determine the number of CFU.

In order to determine the number of invaded bacteria in macrophages, after the aforementioned 2 hour incubation period each well of parallel infected cells were washed with 2x with cell culture grade PBS containing calcium (Ca++) and magnesium (Mg++) (+/+ PBS). RPMI media was then replaced in each well along with 1.0 mg/mL kanamycin. The resultant mixture was incubated for another 2 hours under similar conditions and washed 3x with +/+ PBS. To lyse the macrophages, 1 % Triton X-100 lysis buffer was similarly added to each well. The plate was then shaken for 15 minutes to detach cells, and the resultant bacteria were diluted, plated on TSA, incubated for 1-2 days, and counted.

### Creation of lacZ mutant *B. dolosa* strains

To differentiate between O-antigen mutant and intact strains in competition experiments, a lacZ cassette was inserted one strain of several pairs of near isogenic *B. dolosa* isolates: R-317 and Q-103, R-211 and R-107, R-311 and R-207. All lacZ cassettes were inserted into the O-antigen-intact strain (Q-103, R-107, and R-207). An inverse, R-317 lacZ and Q-103 strain pair was engineered to ensure the lacZ insertion did not impart any costly fitness effects.

Strain R-317 is near isogenic to Q-103, differing by a Y438* SNV in *wbpW* (BDAG_02323/AK34_RS24410), a “mannose-1-phosphate guanylyltransferase” projected to impact O-antigen expression, an intergenic G→T SNV at position 1284152 on chromosome 2, and a synonymous SNV in BDAG_01365 (AK34_RS18940), a “porin.” Strain R-211 differs from R-107 by a P97R SNV in *gmd* (BDAG_02320/AK34_RS24390), a “GDP-mannose 4,6-dehydratase” projected to impact O-antigen expression and no other known SNVs. Strain R-311 differs from R-207 differ by a +G insertion in *wbiH* (BDAG_02309/AK34_RS24330) projected to impact the O-antigen, and S130A in a hypothetical protein AK34_RS02570 (no BDAG identification number).

To constitutively express lacZ, mutant *B. dolosa* strains were generated using pCElacZ^58^ reporter, which utilizes a mini-Tn7 based transposon for stable integration into the chromosome without the need for antibiotic selection. This reporter plasmid was conjugated into each *B. dolosa* strain with the helper plasmid pRK2013 and integration-helped plasmid pTNS3 as described by Choi et al^59^. Conjugates were selected by plating the colonies on LB agar containing trimethoprim (1 mg/ml) and gentamicin (50 μg/ml). Insertions into the *att*Tn7 site downstream of BDAG_04221 (AK34_RS08675) were confirmed by PCR (Forward-PTn7L-ATTAGCTTACGACGCTACACCC and Reverse-bdag-4221 GCGTTCTTGCACCGAACATG).

### Modeling spatial spread of *B. dolosa* via intranasal murine infection

We implemented the murine infection model described in Schaefers et al.^19^ to compare how intact (“medium” phenotype) and disrupted O-antigen phenotypes impact infection establishment. All animal experiments were approved by the Boston Children’s Hospital Institutional Animal Care and Use Committee under assurance number A3303-01 and protocol number 1241. All protocols are compliant with the NIH Office of Laboratory Animal Welfare, the Guide for the Care and Use of Laboratory Animals, the US Animal Welfare Act, and the PHS Policy on Humane Care and Use of Laboratory Animals.

To prepare *B. dolosa* inoculum, overnight cultures of lacZ-marked intact and disrupted O-antigen strains were grown in TSB. After subculturing each strain to log-phase, each culture was diluted with PBS to a final concentration of 2×10^6^ CFU/ml. Before inoculation, samples were plated on PC Agar (*Pseudomonas cepacia* agar, BD biosciences) containing 100 mg/ml of X-gal (5-Bromo-4-chloro-3-indolyl β-D-galactopyranoside) (Thermofisher) to confirm CFU/ml counts.

Mice were anesthetized with ketamine (66.6 mg/kg) and xylazine (13.3 mg/kg) given intraperitoneally. To infect mice with a 50:50 mixture of *B. dolosa* phenotypes, 10 μl of inoculum was inserted into each nostril of mice held in dorsal recumbency. After 7 days, mice were euthanized by CO_2_ overdose, and the lungs and spleens were then aseptically removed. Each organ was weighted, placed into 1 ml of 1% proteose peptone in water, homogenized, and serially diluted and plated on custom PC-X-gal agar. After 36 hours, blue and white colonies were counted to determine the phenotype ratio in each lung. This experiment was performed twice for each of the three strain pairs. We additionally competed the pair Q-103 and R-317 lacZ and pair Q-103 lacZ and R-317 against each other once to account for potential effects of the lacZ cassette (**Fig. S12**).

## Data and code availability

All code used in this project is published under https://github.com/ajporet/b_dolosa_evolution. Sequences from *B. dolosa* samples obtained in this study are uploaded under BioProject PRJNA1063312.

## Contributions

T.D.L. and G.P.P. conceptualized, designed, and jointly supervised the work. G.P.P., T.D.L., S.O.V. collected autopsy samples. A.J.P., C.M., G.K.L. prepared isolates and sequencing libraries. A.J.P., C.M., M.M.S, K.E.M, A.R.C., J.B.G conducted phenotypic assays. A.J.P., M.M.S, K.E.M generated mutant constructs and conducted in vivo experiments. A.Z.U. assisted with patient recruitment and clinical descriptions. A.J.P., T.D.L., M.M.S., and G.P.P. conducted all analyses and interpreted results. A.J.P. and T.D.L. wrote the manuscript with M.M.S and G.P.P. All authors provided feedback on the manuscript.

## Acknowledgments

We thank Robert Fowler, Jonathan Greenberg, Elizabeth Carpino, and Craig Gerard for assistance in obtaining samples. We thank Chris Mancuso, Laura Markey for feedback on the manuscript, and all members of Lieberman lab for advice and support. This work was funded in part by the Richard A. and Susan F. Smith President’s Innovation Award (to G.P.P.) and by funds for the Translational Research for Infection Prevention in Pediatric Anesthesia and Critical Care (TRIPPACC), Program from the Department of Anesthesiology, Critical Care and Pain Medicine at Boston Children’s Hospital (to G.P.P.), and U.S. National Institutes of Health grants DP2-GM140922 (to T.D.L.), and the Burroughs Wellcome Fund via a Career Award at the Scientific Interface, Broad Institute Startup Funds, an Emerging Technologies Opportunity Program award from the Department of Energy Joint Genome Institute, and the National Institute for Allergy and Infectious Disease (AI121932) (to P.C.B.).

