## Supplemental Figures 1-13 for "*De novo* mutations mediate phenotypic switching in an opportunistic human lung pathogen"

### Supplementary Figures

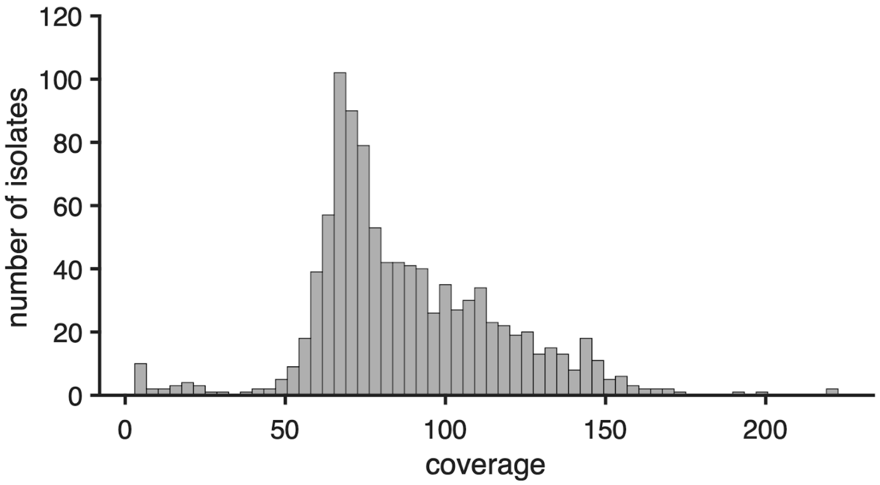


**Supp. Figure 1: 987 *B. dolosa* isolates were sequenced at an average depth of 88x**
A histogram catalogs the coverage of every isolate sequenced for this project. This dataset was sequenced to a mean depth of 87.7x with a standard deviation of 29.3x.

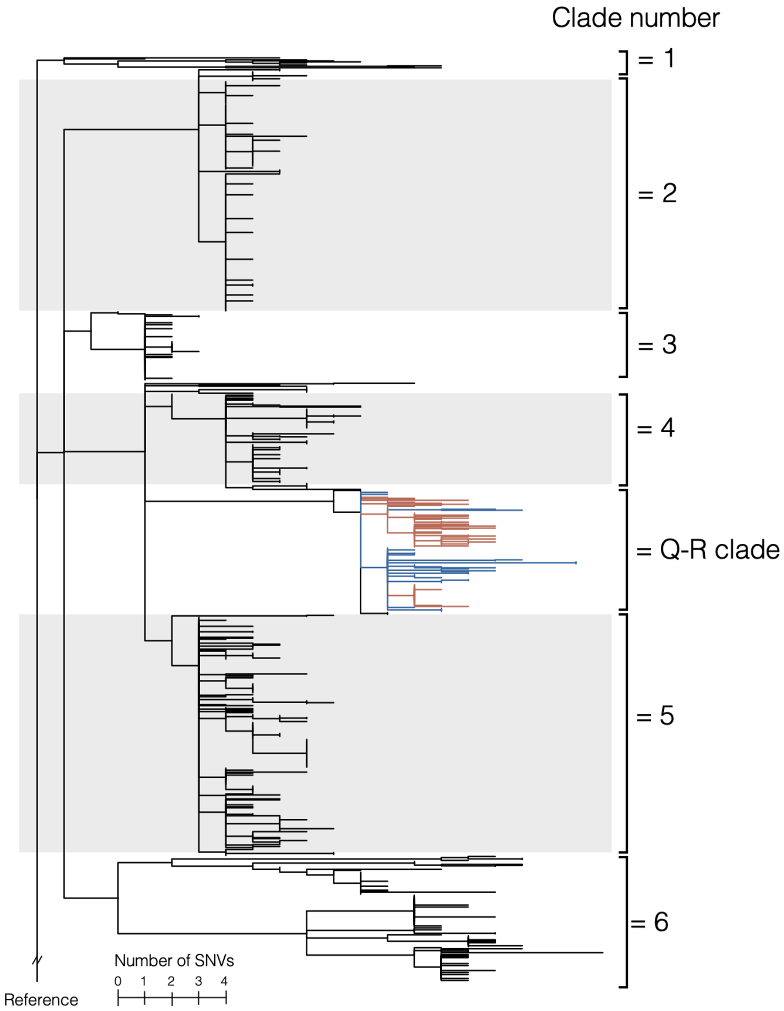


**Supp. Figure 2: A SNV phylogeny of Patient J's, Q's, & R's *B. dolosa* isolates displays transmission patterns.** More detailed version of the subclade of Fig. 2a shaded in gray. We created a maximum parsimony SNV phylogeny of all sequenced isolates from Patients J, Q, and R. Branch colors indicate patient, with Patient Q marked as blue, Patient R as red, and Patient J as black. Six major clades are observed within J's isolates. These are numbered for reference in other supplementary materials. Eleven samples share 3 SNVs common to clades 4, 5, and Q-R but lack shared SNVs that define each of these clades; these samples are therefore not included in any numbered clade.

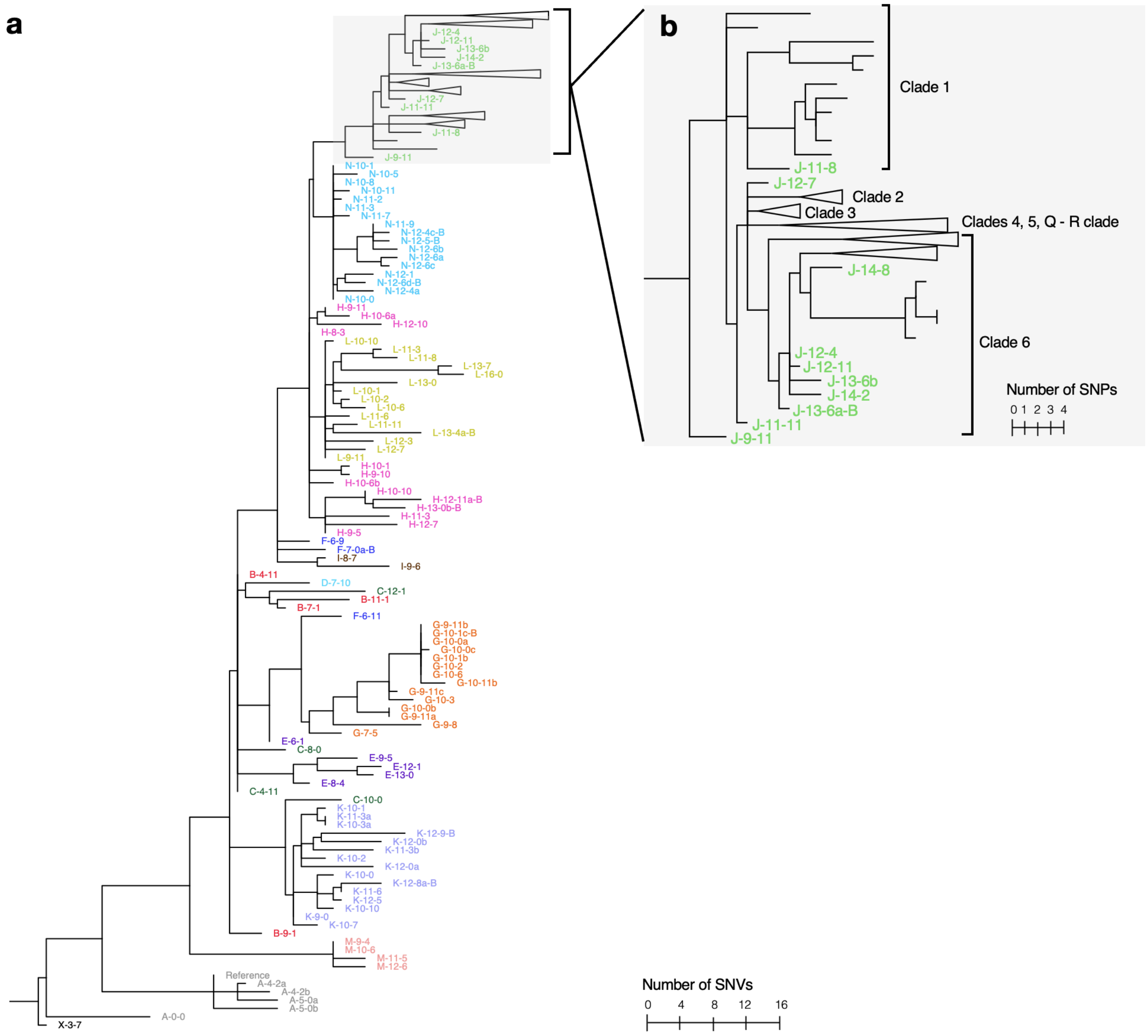


**Supp. Figure 3: Comparing Patient J's *B. dolosa* isolates to those from a prior *B. dolosa* outbreak** (a) Same tree as in Fig. 2a, with isolates from the Lieberman and Michel et al.[15] study expanded and Patient J's isolates compressed. Isolates collected in the prior outbreak are colored by patient. We also included an outgroup isolate X-3-7 (in black). Isolates from Lieberman and Michel et al. are named according to patient and time as previously (ex. C-14-11 was recovered from Patient C, 14 years and 11 months after isolation of the first outbreak strain). Samples with a "-B" at the end of their label indicate a blood source; otherwise, all remaining samples profiled in 2011 were collected from sputum. Isolates with the same subject ID and date are distinguished by letters (ex. J-13-6a-B and J-13-6b). (b) A zoom in on Patient J's samples from Lieberman and Michel et al.'s[15] investigation enables positioning of new samples in context to those obtained a decade prior. Clade names correspond to the tree in Supp. Fig. 2.

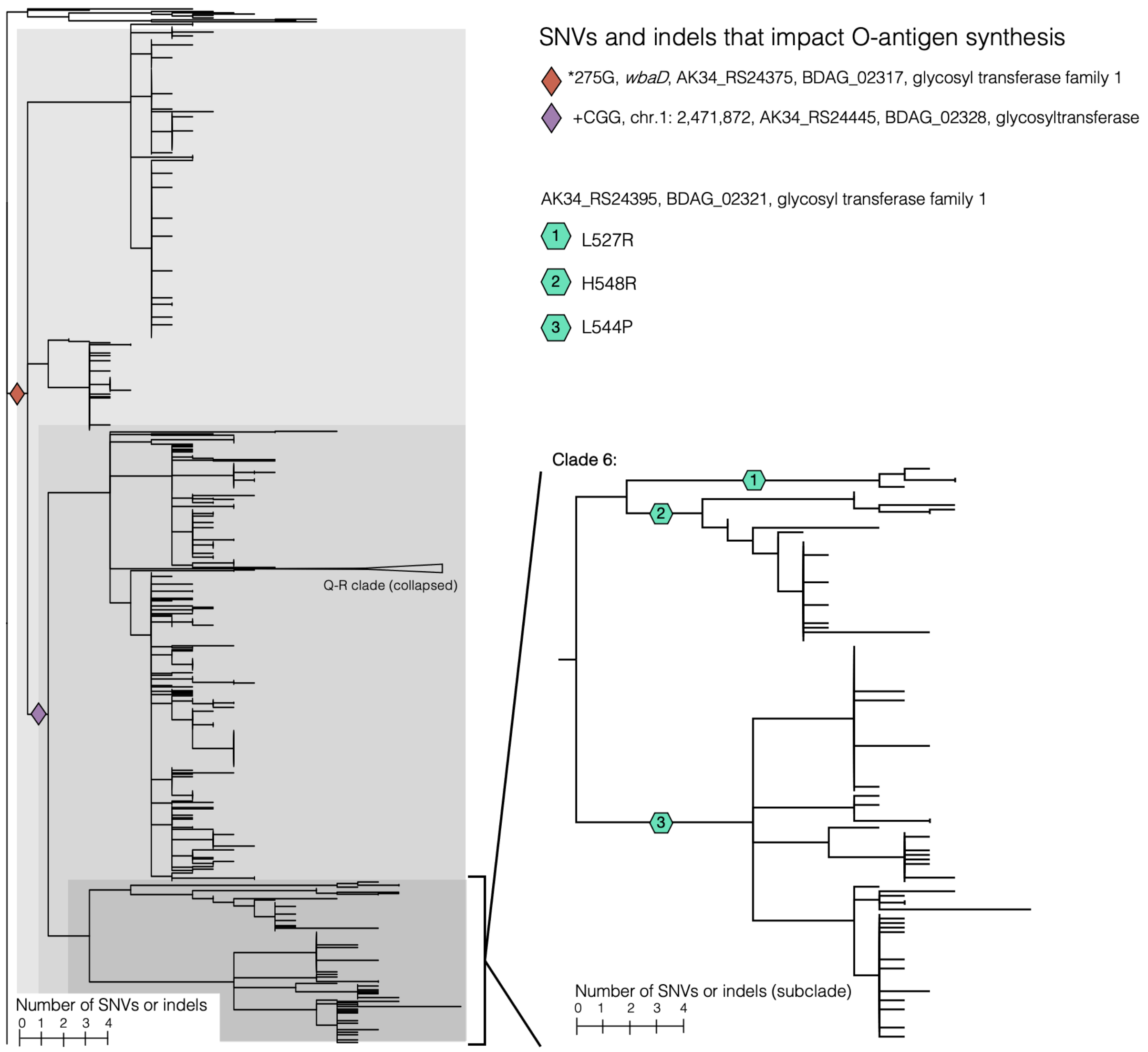


**Supp. Figure 4: Over long-term infection, *B. dolosa* accumulates multiple SNVs and indels that impact O-antigen synthesis.** A maximum parsimony phylogeny of isolates from Patients J, R, and Q generated from both SNVs and O-antigen-affecting indels, where each indel is represented as a single, independent mutation event (Methods). Diamonds and hexagons demark the emergence of O-antigen-affecting mutations. Clade Q-R is collapsed for simplicity, and a zoom-in on clade 6 reveals clustered O-antigen-affecting mutations. Of 809 strains acquired from Patient J's autopsy, 11 (1.4%, clade 1, white) contain a stop codon, while 317 (39%, clades 2 & 3, light gray) contain only a *wbaD* (BDAG_02317/AK34_RS24375) stop codon reversion. A total of 355 (44%, clades 4, 5, 11 ungrouped samples, and 4 isolates within the Q-R clade, medium gray) contain a further +CGG insertion in BDAG_02328 (AK34_RS24445), and 126 (16%, clade 6, dark gray) contain one of three additional SNVs in BDAG_02321 (AK34_RS24395).

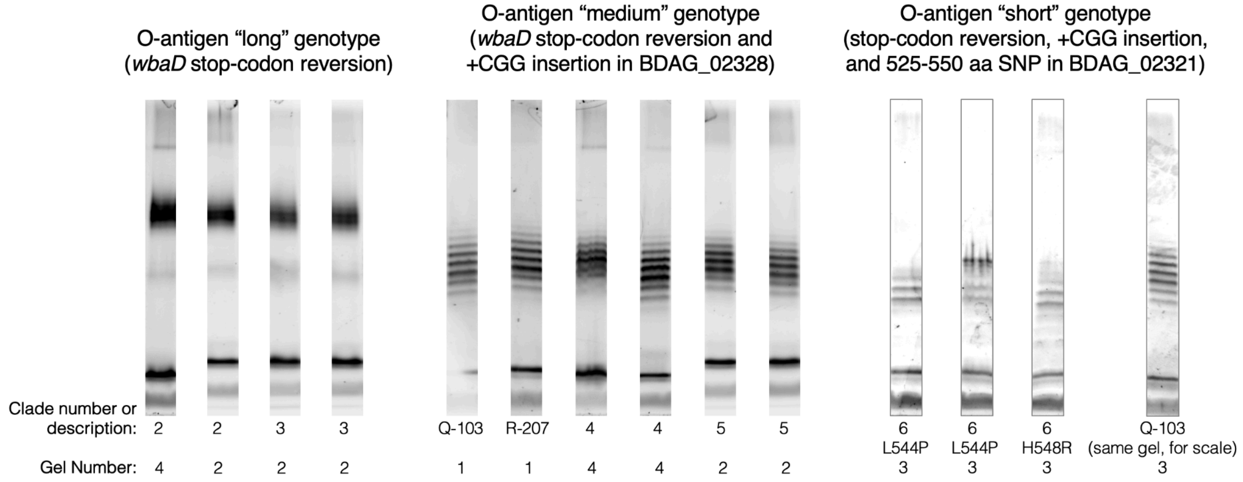


**Supp. Figure 5: Different O-antigen phenotypes coexist in *B. dolosa* lung infection.** O-antigen phenotypes of diverse *B. dolosa* clones from across Fig. S4's phylogeny. Strains containing only a *wbaD* stop codon reversion (clades 2 and 3) display a "long" banding pattern. Strains containing a +CGG insertion in BDAG_02328 (AK34_RS24445) present a "medium" banding pattern (clades 4,5, Q-R, and 11 ungrouped samples); this is the phenotype that is inferred to have first infected Patients Q and R. Isolates from clade 6 display acquired one of three mutations in the 525-550 amino acid region of BDAG_02321 (AK34_RS24395). Phenotypes of both BDAG_02321 L544P and H548R display a "short" banding pattern; the third independent BDAG_02321 mutation (L527R) was not phenotyped. To contextualize the "short" banding length, a "medium" length isolate (Q-103) imaged on the same gel is shown to the right. Two "standard" isolates, Q-103 and R-221, were imaged on all gels to aid in banding length comparison; raw gel images with these "standard" isolates can be found under the Github repo: ajporet/b_dolosa_evolution. Gel numbers correspond to file names in Github.

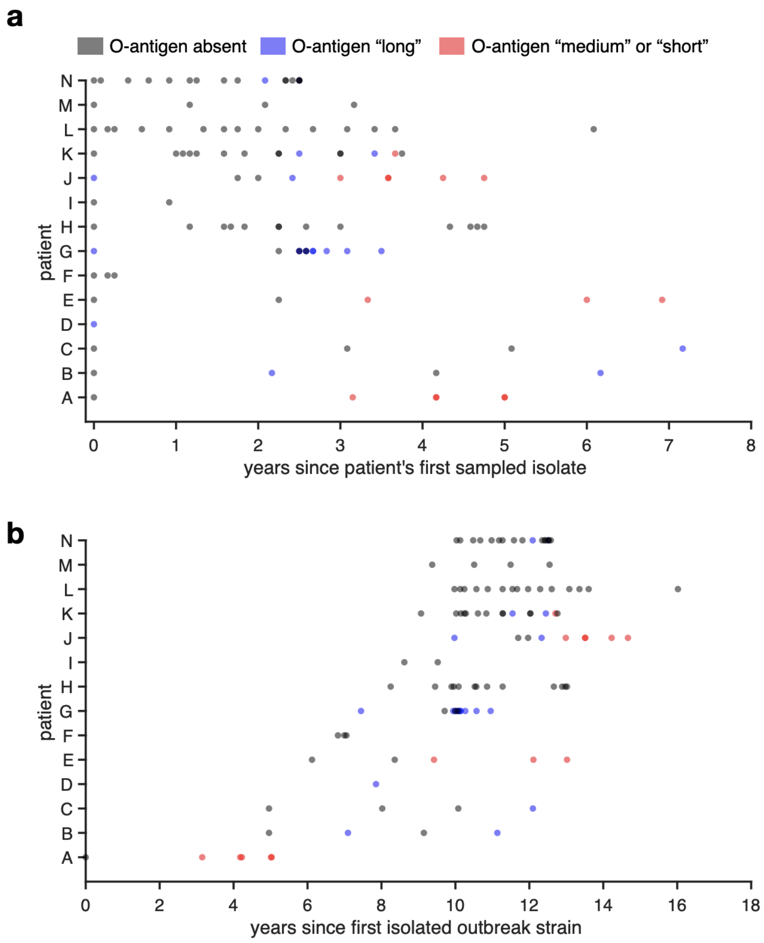


**Supp. Figure 6: Comparing sampling date and time to O-antigen genotype reveals strain coexistence.** (a) O-antigen banding patterns are predicted for 112 previously-published *B. dolosa* genomes[15] collected during a historical outbreak. Each isolate is represented by a single, shaded dot; the color of each data point represents its predicted O-antigen phenotype. Isolates that contain a *wbaD* stop codon are classified as “absent " (black). Those with a stop codon reversion are marked as “long” (blue), and those with an additional mutation in a gene predicted to affect O-antigen function are categorized as “medium or short” (red). To understand how *B. dolosa*’s O-antigen evolves over the course of chronic infection, we plotted the sampling date of each isolate relative to the time of each patient’s first sequenced *B. dolosa* isolate. In 5 out of 14 patients (N, K, J, G, B) ancestral, O-antigen-absent strains are recovered years after an O-antigen-expressing (long or medium/short) strain, highlighting co-existence of phenotypes. Additionally, four patients (K, J, E, and A) contain isolates with additional mutations predicted to shorten the O-antigen (red, Fig. S7). **(b)** The same data plotted relative to sampling date.

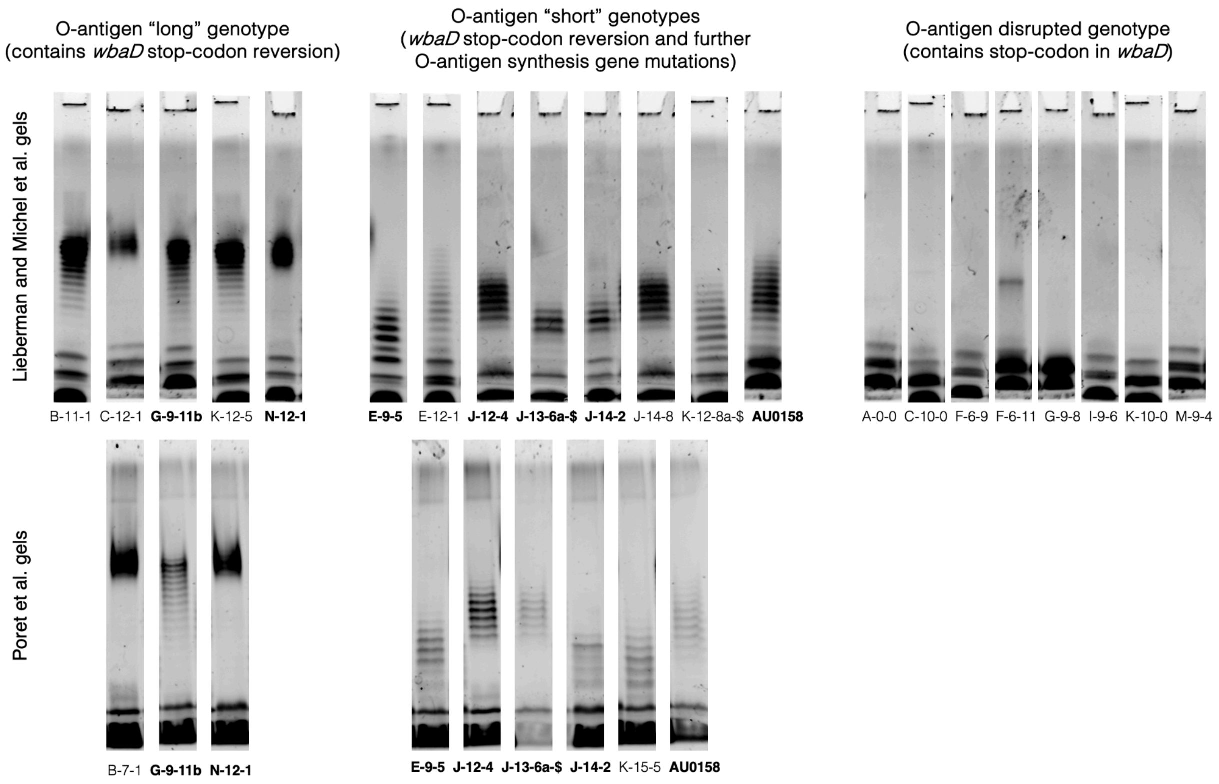


**Supp. Figure 7: Reanalysis of a prior *B. dolosa* outbreak reveals O-antigen modification beyond an initial gain-of-function mutation.** O-antigen banding patterns of *B. dolosa* isolates from a historical 14-person outbreak. O-antigen phenotypes produced by Lieberman and Michel et al.[15] can be grouped into three classes with an associated genotypic signature. "Disrupted" banding patterns contain a stop codon in gene *wbaD* and "long" banding patterns revert this stop codon. All isolates that acquire a further mutation in an O-antigen-affecting gene display a "short" banding pattern. The exact banding length of "short" isolates varies depending on the O-antigen-affecting mutation and genotypic background. To ensure compatibility between newly created gels and those published by Lieberman and Michel et al.[15], previously-published isolates were re-phenotyped. Bolded sample names indicate an isolate is present in both datasets. Banding patterns appear the same between both analyses.

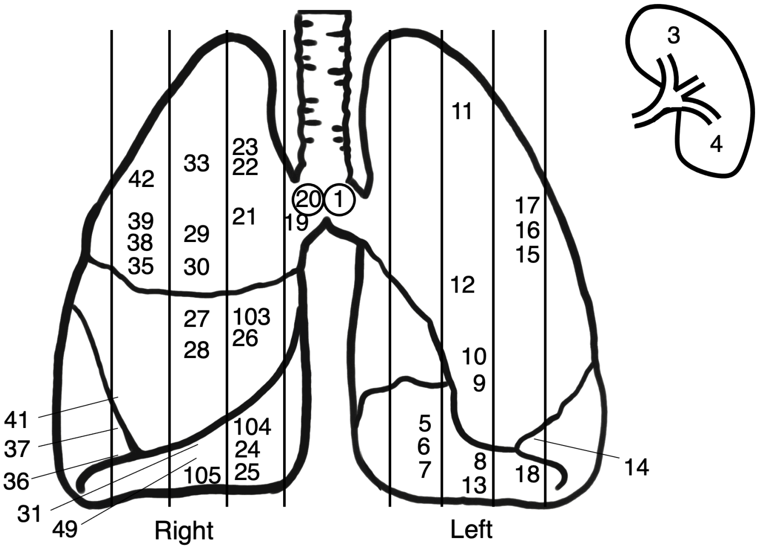


**Supp. Figure 8: Diagram of lung sites sampled in Patient J (autopsy).** The anatomical location of each sampled lung site is diagrammed. Vertical lines indicate dissection cut sites. Circled numbers indicate that sampled tissue is from a mediastinal lymph node. Samples 3 and 4 were taken from splenic tissue. Lung figure is modified from a diagram by Patrick J. Lynch and C. Carl Jaffe (goo.gl/iC8AjM), CC-BY-2.5.

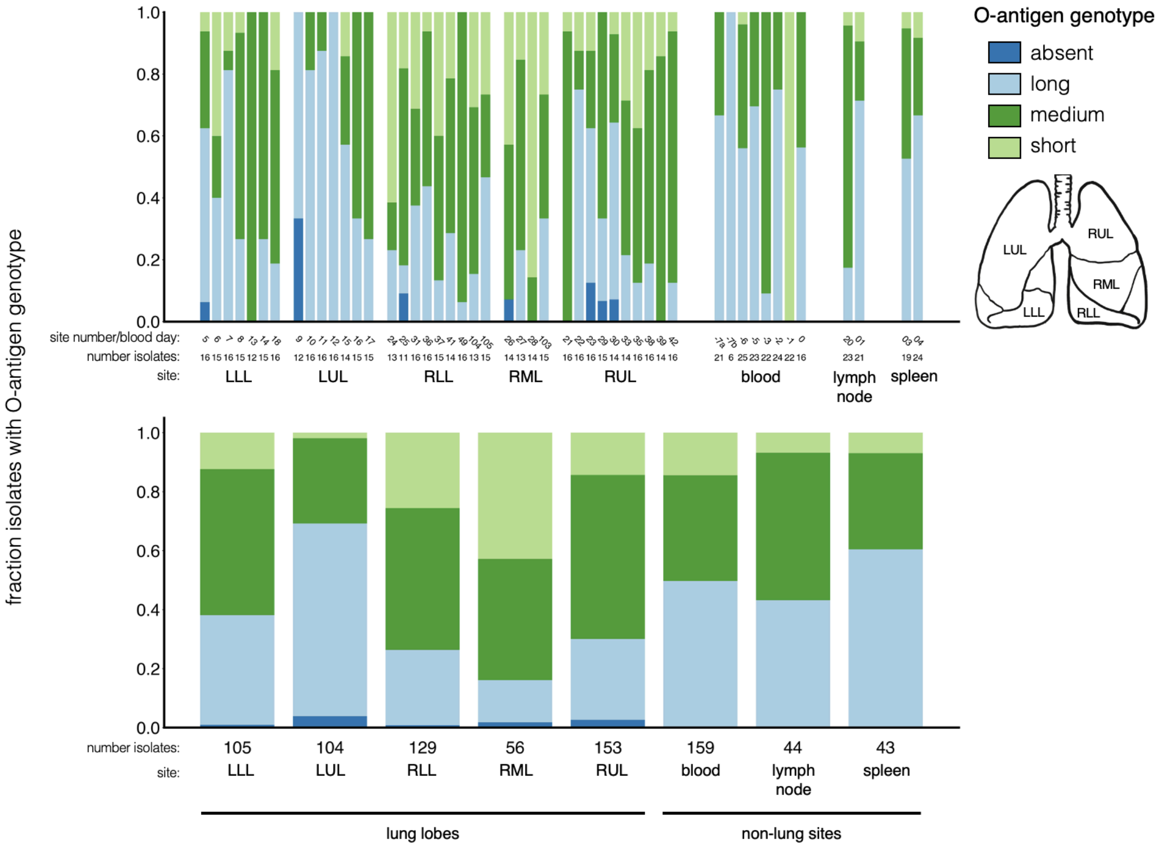


**Supp. Figure 9: Distribution of O-antigen genotypes across lung lobes.** The distribution of O-antigen genotype classes (as defined in main text and Fig. S4, S5) is grouped by anatomical site. Lung, spleen, and lymph node isolates are stratified by sampling site. Blood culture isolates are separated by day of acquisition with 0 marking the day of death. A week prior to death, two independent blood samples were cultured; these are marked as "-7a" and "-7b." Ancestral, O-antigen-absent genotypes are found in all 5 lung lobes. Modest differences in long, medium, and short genotype abundance are observed among lobes ($P < 10^{-10}$; Chi square test).

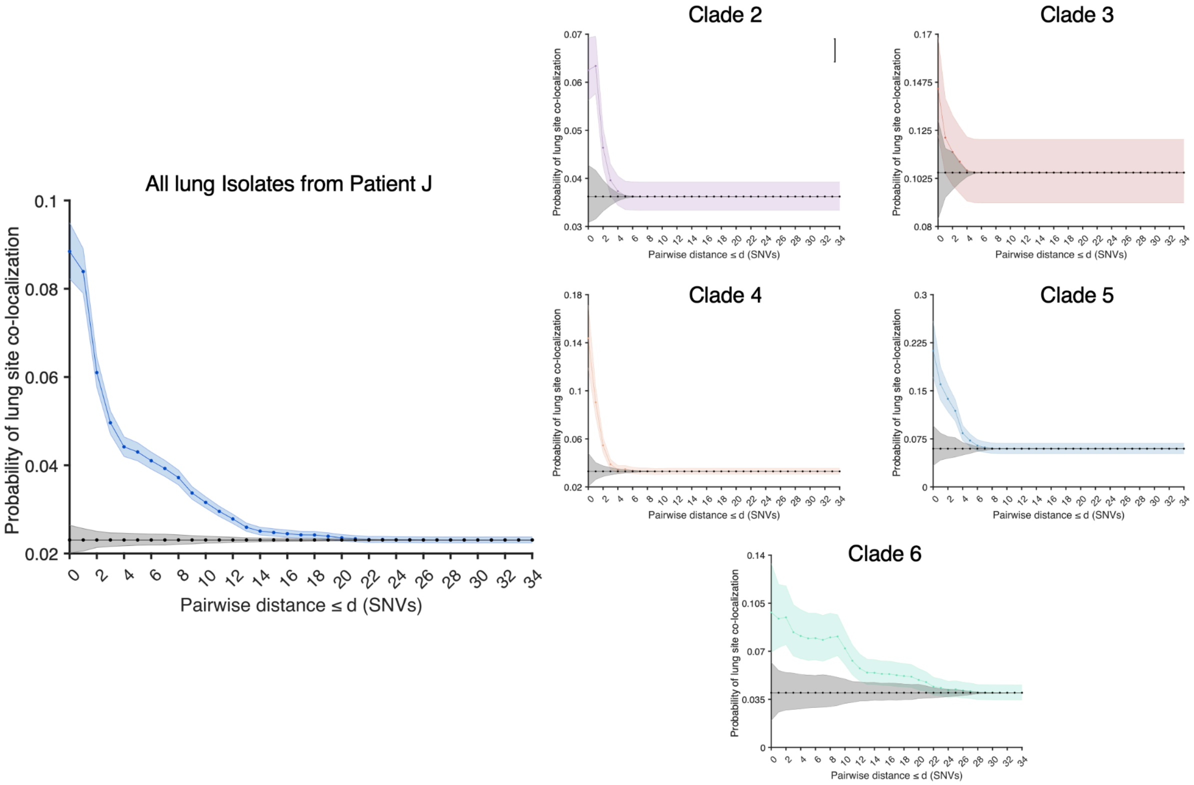


**Supp. Figure 10: Weak spatial segregation of *B. dolosa* genotypes is observed within a CF lung autopsy.** Using the methods of Chung et al.[21], we examined whether *B. dolosa* genotypes within Patient J's lung are spatially segregated; i.e, that the same genotype is likely to cluster within the same lung site. We calculated the probability that isolate pairs separated by d or less SNVs are located in the same sampling site compared to a null model (1000 trials of randomly shuffling lung sites). The black line indicates the null model, and error bars indicate a 95% confidence interval. We calculated the probability of lung-site colocalization for all of Patient J's isolates and each major clade to account for niche evolution that could render one clade spatially diffuse or concentrated. Compared to Chung et al., only minimal spatial correlation between genotype and lung site is observed using these models with the exception of clade 6.

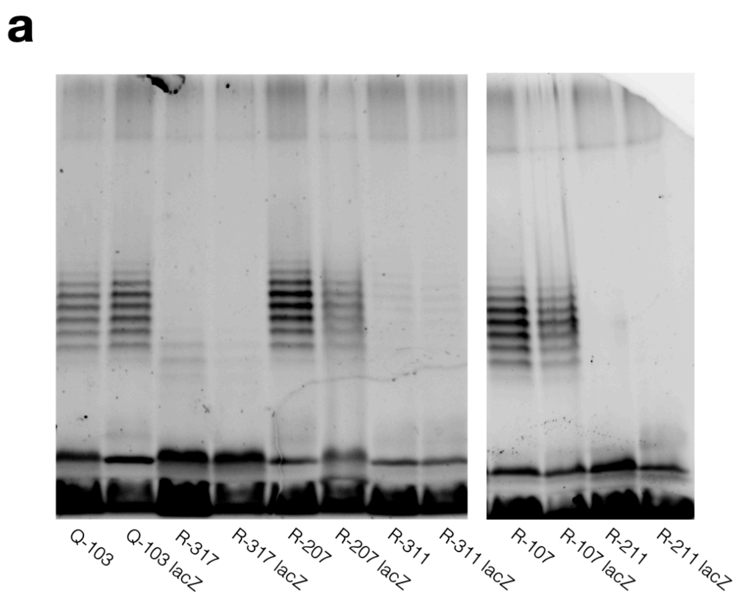


**Supp. Figure 11: Lipopolysaccharide (LPS) staining shows O-antigen differences between strains used in murine model and macrophage assays.** (a) Each pair of *B. dolosa* strains used in the murine infection model were extracted and imaged on the same gel. O-antigen staining patterns show no obvious differences between lacZ marked and unmarked strains. (b) All strains used in the Fig. S13b macrophage invasion experiments were imaged on the same gel, displaying the expected O-antigen phenotype.

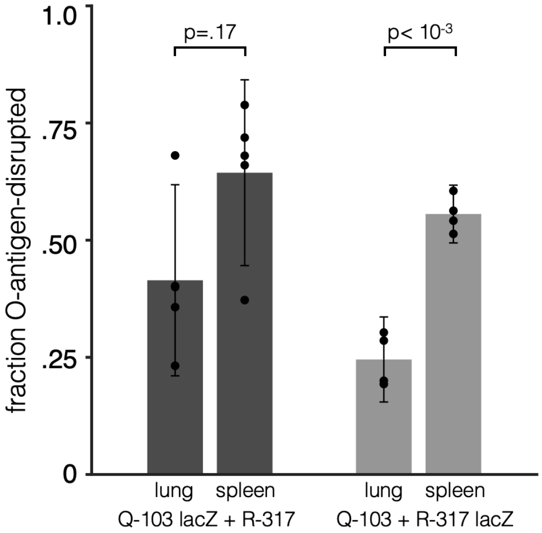


**Supp. Figure 12: LacZ cassette insertion into *B. dolosa* does not impact a murine infection model outcome.** To ensure the LacZ cassette insertion used to differentiate between O-antigen mutant and wild type strains did not bias strain fitness, we conducted a label swap in which a LacZ cassette was integrated into both Q-103 and R-317 strains separately. Experiments were performed with both labeled pairs on the same day. Competitions of Q-103 lacZ vs R-317 and Q-103 vs R-317 lacZ showed similar results to those in Fig. 4a (paired T-test, Q-103 lacZ-R-317: P= .17, Q-103 - R-317 lacZ: $P < 10^{-3}$). Outlier points seen in the Q-103 lacZ + R-317 lung and spleen occur within the same mouse. Data presented here represent a distinct replicate from that shown in Figure 4a; results in Figure 4a come from 2 experimental batches containing all 3 strain pairs shown.

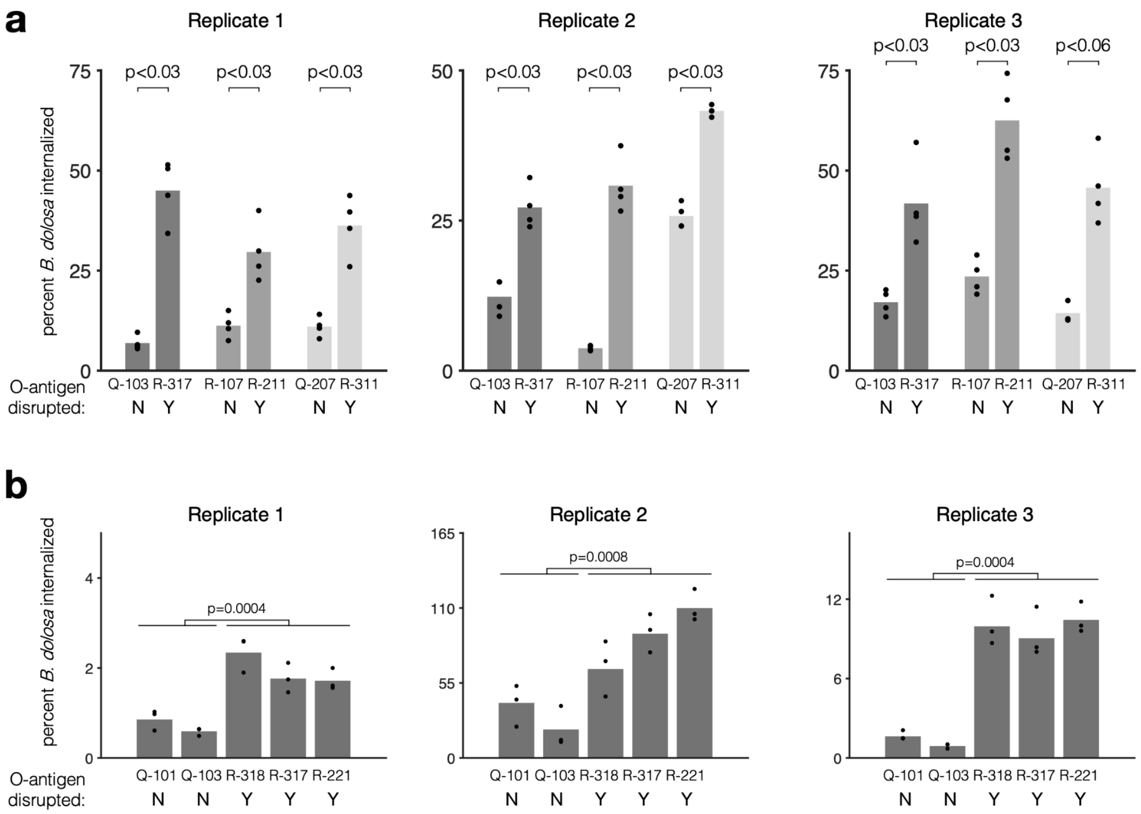


**Supp. Figure 13: Kanamycin exclusion assays demonstrate that O-antigen disruption increases *B. dolosa* invasion/survival within macrophages.** Macrophages were infected for 2 hours with near-isogenic *B. dolosa* strains with and without an O-antigen-disrupting mutation and then incubated for 2 hours with kanamycin, which kills extracellular bacteria. The number of intracellular bacteria is compared to the total bacteria obtained from an identical culture without kanamycin treatment. (a) All *B. dolosa* strain pairs used in the murine experiments were compared against each other, demonstrating significant differences in macrophage invasion (P < .07, Wilcoxon rank sum test, p-values are uncorrected). Replicate 1 is also shown in Fig. 4b. (b) A separate panel including two O-antigen wild-type (Q-103, Q-101) and mutant (R-318, R-317, R-221) strains display the same expected phenotype in the macrophage assays as shown in Fig. 4b. All measurements of O-antigen mutant strains are aggregated and compared to O-antigen wild-type strains (P < .001, Wilcoxon rank sum test, p-values are uncorrected).
